# Uncovering the hidden structure of dynamic T cell composition in peripheral blood during cancer immunotherapy: a topic modeling approach

**DOI:** 10.1101/2023.04.24.538095

**Authors:** Xiyu Peng, Jasme Lee, Matthew Adamow, Colleen Maher, Michael A. Postow, Margaret K. Callahan, Katherine S. Panageas, Ronglai Shen

**Author notes:** **Contributions** X.P. contributed to the original draft and performed bioinformatics data analysis. X.P., R.S., K.S.P. conceived and designed the algorithm. J.L. contributed to the pre-gating analysis of flow cytometry data. R.S., K.S.P., and M.K.C. developed the initial study concept and oversaw all data generation and analysis. C.M. extracted and analyzed clinical data. M.A. generated and analyzed flow cytometry data. M.A.P. contributed to the initial concept and to the acquisition and analysis of clinical data. X.P, J.L., R.S., K.S.P., and M.K.C reviewed and edited the manuscript. All authors reviewed and approved the final manuscript.

## Abstract

Immune checkpoint inhibitors (ICIs), now mainstays in the treatment of cancer treatment, show great potential but only benefit a subset of patients. A more complete understanding of the immunological mechanisms and pharmacodynamics of ICI in cancer patients will help identify the patients most likely to benefit and will generate knowledge for the development of next-generation ICI regimens. We set out to interrogate the early temporal evolution of T cell populations from longitudinal single-cell flow cytometry data. We developed an innovative statistical and computational approach using a Latent Dirichlet Allocation (LDA) model that extends the concept of topic modeling used in text mining. This powerful unsupervised learning tool allows us to discover compositional topics within immune cell populations that have distinct functional and differentiation states and are biologically and clinically relevant. To illustrate the model’s utility, we analyzed ∼17 million T cells obtained from 138 pre- and on-treatment peripheral blood samples from a cohort of melanoma patients treated with ICIs. We identified three latent dynamic topics: a T-cell exhaustion topic that recapitulates a LAG3+ predominant patient subgroup with poor clinical outcome; a naive topic that shows association with immune-related toxicity; and an immune activation topic that emerges upon ICI treatment. We identified that a patient subgroup with a high baseline of the naïve topic has a higher toxicity grade. While the current application is demonstrated using flow cytometry data, our approach has broader utility and creates a new direction for translating single-cell data into biological and clinical insights.

## Introduction

Cancer immunotherapies with immune checkpoint inhibitors (ICIs) are revolutionizing cancer treatment^1^. ICIs, given as monotherapy or in combination, have proven efficacious in multiple types of cancer and it is estimated that approximately 44% of cancer patients in the United States are eligible to receive ICIs^2^. However, patient tumor response and toxicity under different treatment regimens are highly heterogeneous. Patients with melanoma who receive CTLA-4 and PD-1 combination blockade have a higher response rate but are more likely to experience immune-related adverse events (irAEs) compared to monotherapy^3–5^. Thus, it is crucial to gain a deeper understanding of the immune mechanisms and pharmacodynamics of ICIs to personalize treatment options, and improve therapeutic benefit while minimizing toxicity for patients^6^.

Flow cytometry analysis has become an important tool to study tumor microenvironment as well as patients’ peripheral blood samples in the context of immunotherapy. Several biomarkers examining functional cell types have been identified to predict treatment response or define resistance mechanisms to ICIs^7–9^. These analyses commonly focus on a limited number of pre-specified cell types determined from prior domain knowledge, potentially overlooking important unmined subpopulations. Furthermore, recent advances in flow and mass cytometry have significantly improved the throughput allowing 30-50 markers measured simultaneously at single-cell resolution^10^, that allows for the exploration of a much larger number of possible cell subsets. Such high-parameter flow cytometry data when performed on longitudinally collected samples are exceedingly complex and pose a great analytical challenge to delineate cell type composition from millions of single cells and map the temporal evolution of cell types over time. Sophisticated statistical and computational tools are needed to fully leverage the complexity and richness of high-parameter single-cell data in order to expedite biomarker discovery in cancer immunotherapy.

In recent years, there have been consorted efforts to advance the development of cutting-edge computational methods for flow cytometry including visualization, clustering, and lineage tracing of cell populations as reviewed in Aghaeepour *et al.*^11^. The current state-of-the-art approach allows refined cell type classification and visualization. However, it remains a challenge to link such output with the clinical outcomes due to the lack of a framework to quantify cell type composition and associated functional states at the individual sample level. In addition, methods to address temporal evolution using flow cytometry data are lacking.

To fill this gap, we present a novel statistical and computational framework that is inspired by works developed in monitoring temporal dynamics of bacterial strains^12,13^. We adapt the Latent Dirichlet Allocation (LDA) model^14^ to investigate the pharmacodynamics of T cell compositions in peripheral blood of ICI-treated cancer patients early after treatment initiation. LDA is a generative statistical model for the identification of hidden structures in large data and is widely applied for topic discovery in text mining analysis. Here we present a novel application of LDA to understand the temporal evolution of T cells in flow cytometry data to track early pharmacodynamic changes after exposure to ICIs (Fig. 1a). In an unsupervised fashion, LDA explores the hidden structure and identifies latent topics with interpretable features relating to biologically relevant function states (Fig. 1b), allowing for the discovery of potential biomarkers of clinical relevance. This approach can be used to predict outcomes and quantify the pharmacodynamics of immunotherapy.

**Fig. 1:**
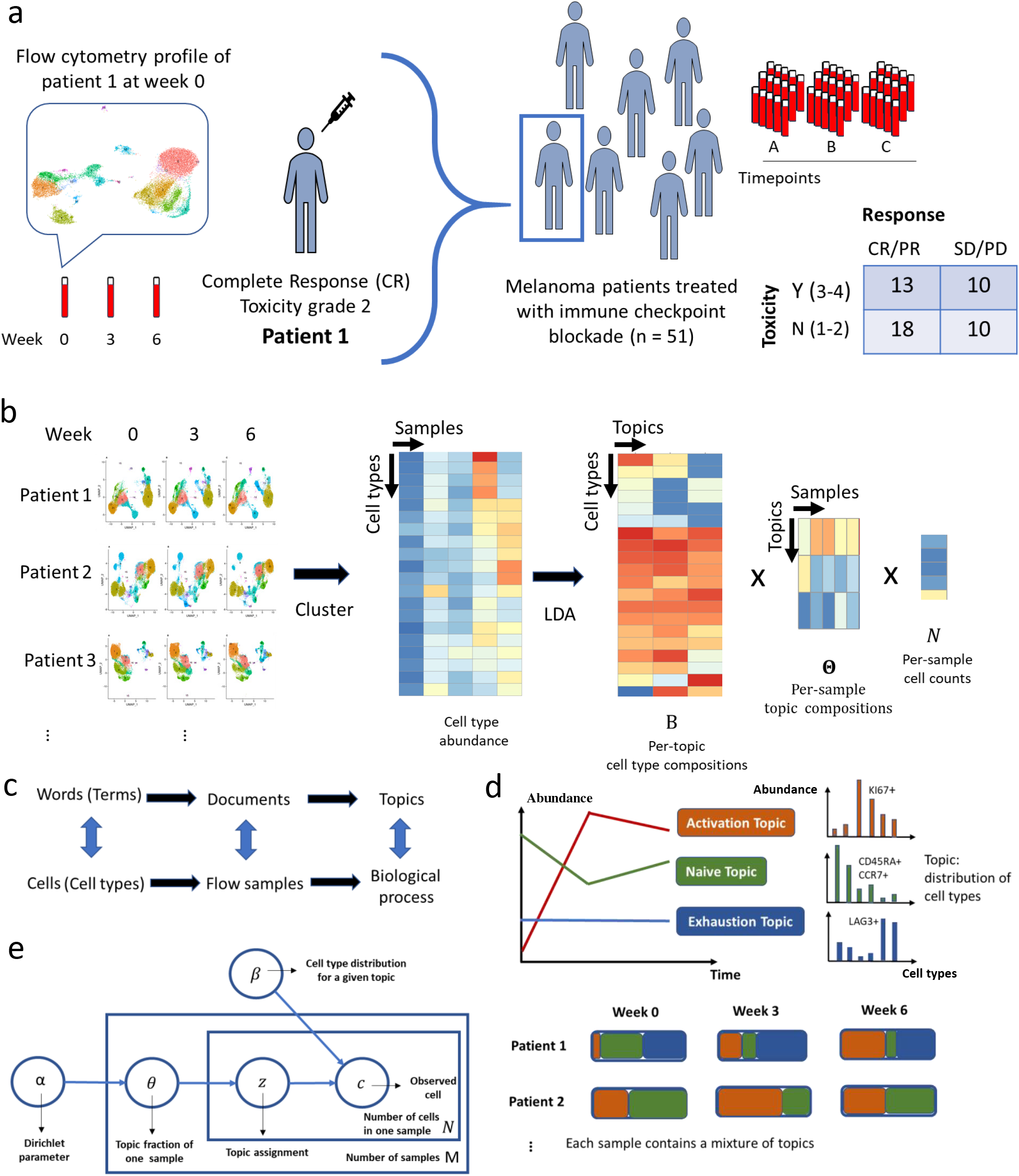
Latent Dirichlet Allocation reveals hidden structures in flow cytometry data. a. Data overview. b. Deconvolution of flow cytometry data with Latent Dirichlet Allocation (LDA) model after pooled clustering analysis. c. The analogy between text analysis and flow cytometry analysis. d. Fractional membership of topics within each sample and its evolution over time. e. Graphic representation of LDA model.

## Results

### Method Overview

We present a topic model approach for mining large-scale high-dimensional flow cytometry data from longitudinally collected patient samples. Motivated by the similarities between text data mining and flow cytometry analysis, LDA considers *cells as words*, *cell types* as *terms*, *patient samples* as *documents*, and *biological processes* as *topics* (Fig. 1c). It assumes each cell in a patient sample arises from a mixture of topics, each of which is distributed over cell types. The cell types can be obtained through a graph-based clustering of single cells from pooled samples (Fig. 1b). Then cell type-by-sample count matrix is decomposed by LDA into three matrices:

1. cell type-by-topic matrix, *B,* for topic content
2. topic-by-sample matrix, Θ, for topic prevalence
3. vector of cell counts *N*.

The cell type-by-topic matrix represents topics as different discrete distributions over cell types, thus facilitating the linkage between topics and cell types. Each topic is a weighted combination of a specific set of cell types that may be functionally related. Within each topic, cell types that show similar abundance patterns across patient samples are likely to be involved in the same biological process. In contrast to the traditional approach of assessing one cell type at a time, LDA provides a unified approach to systematically evaluate all cell types simultaneously and gain insight into the underlying biological processes through their co-occurring patterns.

The topic-by-sample matrix displays topic proportions estimated within each sample. This allows us to characterize and quantify topic composition at the individual sample level and track the topic evolution over time (Fig. 1d). Patients with similar topic composition and temporal dynamics may share the similar clinical outcomes and pharmacodynamic profiles as we will describe in detail in <u>Methods</u>. Below, we illustrate how LDA deconvolutes the longitudinal flow cytometry data to characterize topics with novel biological insights using a data example.

### Data

The large-scale flow cytometry dataset we analyzed contains ∼17 million T cells from a cohort of 51 melanoma patients (138 samples) treated with a combination of anti-CTLA-4 and anti-PD-1 ICI as part of a phase II clinical trial (NCT03122522)^15^. The clinical outcome data (response, overall survival (OS), progression-free survival (PFS), toxicity) of the cohort have been previously reported^15^ and are shown in Supplementary Data File S1. Based on pre-treatment peripheral blood samples, our prior work on a large cohort has classified patients into three ‘immunotypes’ (LAG+/LAG-/PRO) that are correlated to survival and response^16^, which we also include in the analysis. Nearly half of patients (45%) experienced severe (>= grade 3) immune-related adverse events (irAEs) and 61% of patients responded (Complete Response, CR or Partial Response, PR) to the ICI treatment (Fig. 1a). Flow cytometry was performed using an X50 panel that measures 29 markers for each single cell (a complete list of markers described in <u>Methods</u>), including checkpoint blockade biomarkers (e.g. PD1, CTLA4, LAG3) and T cell lineage markers (e.g. CD45RA, CCR7, CD27, CD28). Staining was performed on the cryo-banked peripheral blood mononuclear cells (PBMCs) collected at three time points for each patient: week 0 (pre-treatment), week 3 and 6 (post-treatment).

### Identification of T cell types and composition across patient samples

Before applying the LDA model, we first identified T cell types via the Louvain algorithm, a popular data-driven graph-based clustering method^17^, after pooling viable CD3+ cells from all patient samples at all time points together to allow the comparison of consistent T cell clusters across multiple samples. The optimal clustering resolution was determined based on average Silhouette scores^18^ and manual evaluation (See details in <u>Methods</u>). The 20 main T cell clusters with relative abundance > 0.1% are displayed in the Uniform Manifold Approximation and Projection (UMAP) (Fig. 2a), where CD4 and CD8 T cells are separated into two distinct parts (Fig. 2b). The marker expression profile in the T cell clusters is shown in Fig. 2c. Based on the lineage marker CD45RA and CCR7 (Fig. 2d), we are able to further identify T cell clusters with different differentiation states, including the naïve T cell clusters (Tn, CCR7+CD45RA+), central and effector memory T cell clusters (Tcm, CCR7+CD45RA-, and Tem, CCR7-CD45RA-), and CD45RA+ effector memory T cell clusters (Temra, CCR7-CD45RA+). Moreover, we identified two clusters, one CD4 Tcm cluster (cluster 8) and one CD8 Tem cluster (cluster 12), that highly express KI67, a proliferation marker recognized in previous studies^9^ (Fig. 2d).

**Fig. 2:**
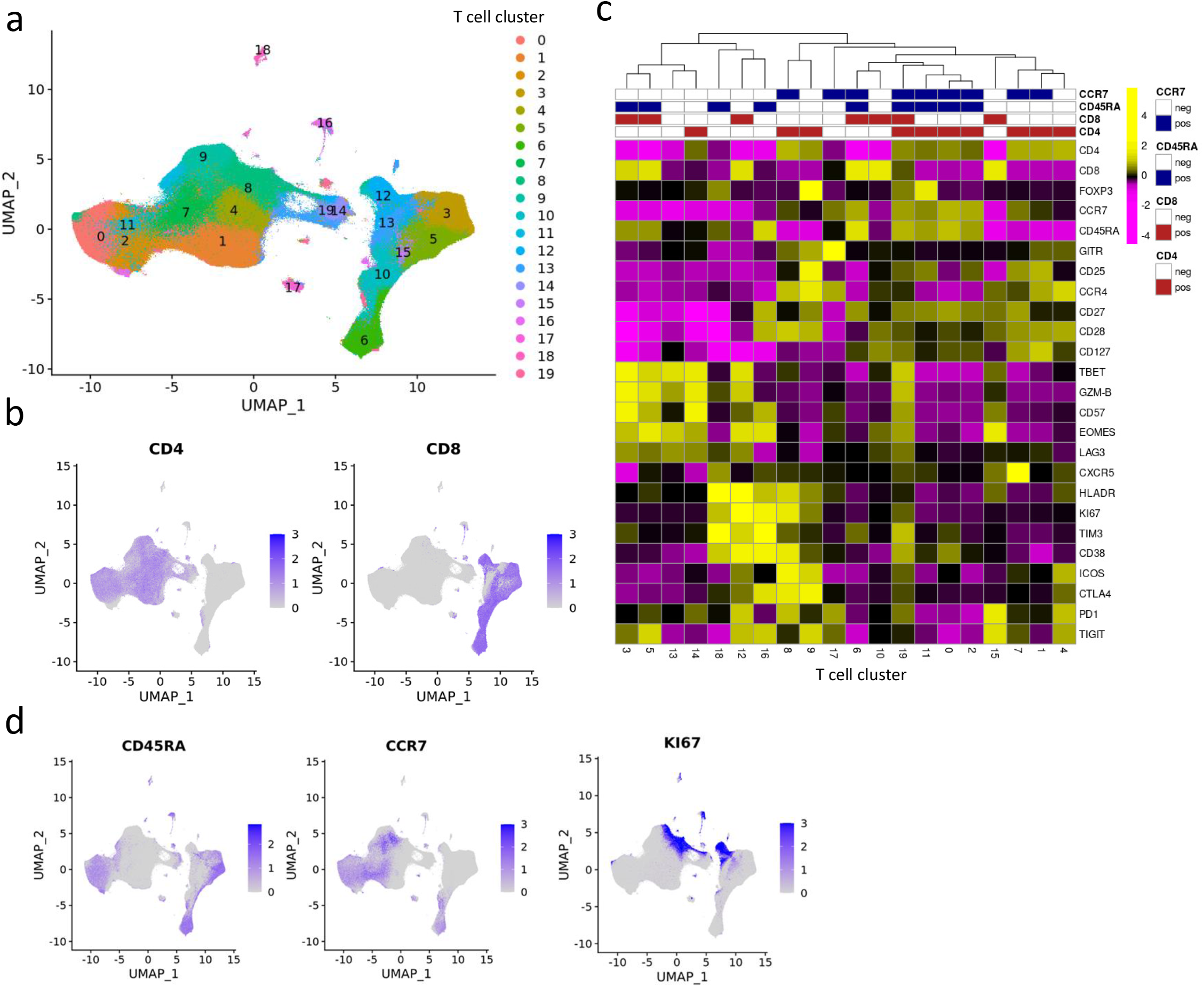
Identification of T cell clusters in the X50 flow cytometry data. a. UMAP plot of T cell clusters. b. UMAP plot of T cells overlaid with the expression of CD4 and CD8. c. Heatmap displaying average marker expression (scaled) of markers in each cluster. d. UMAP plot of T cells overlaid with the expression of CD45RA, CCR7, and KI67.

### Latent Dirichlet Allocation reveals hidden structures in flow cytometry data

The T cell clusters we identified are inter-correlated as governed by the underlying functional and differentiation states. We applied LDA and uncovered K = 3 latent topics, which capture the major patterns underlying the data. The determination of the number of topics K is described in Methods.

We first evaluate each topic by visualizing the weights *β_k_* for every single topic, where a topic is represented as a distinct probability distribution over the T cell clusters (Fig. 3a). Based on the pattern of this distribution we define three topics as **activation** topic, **naïve** topic and **exhaustion** topic based upon domain knowledge. The activation topic is mainly contributed by memory T cell clusters (Tcm/em), and later we will show that these clusters capture the major pattern of T cell expansion after ICI. The naïve topic has high probability weights over the naïve T cell clusters (Tn) while the exhaustion topic consists of exclusively terminally differentiated T cell clusters (Temra).

**Fig. 3:**
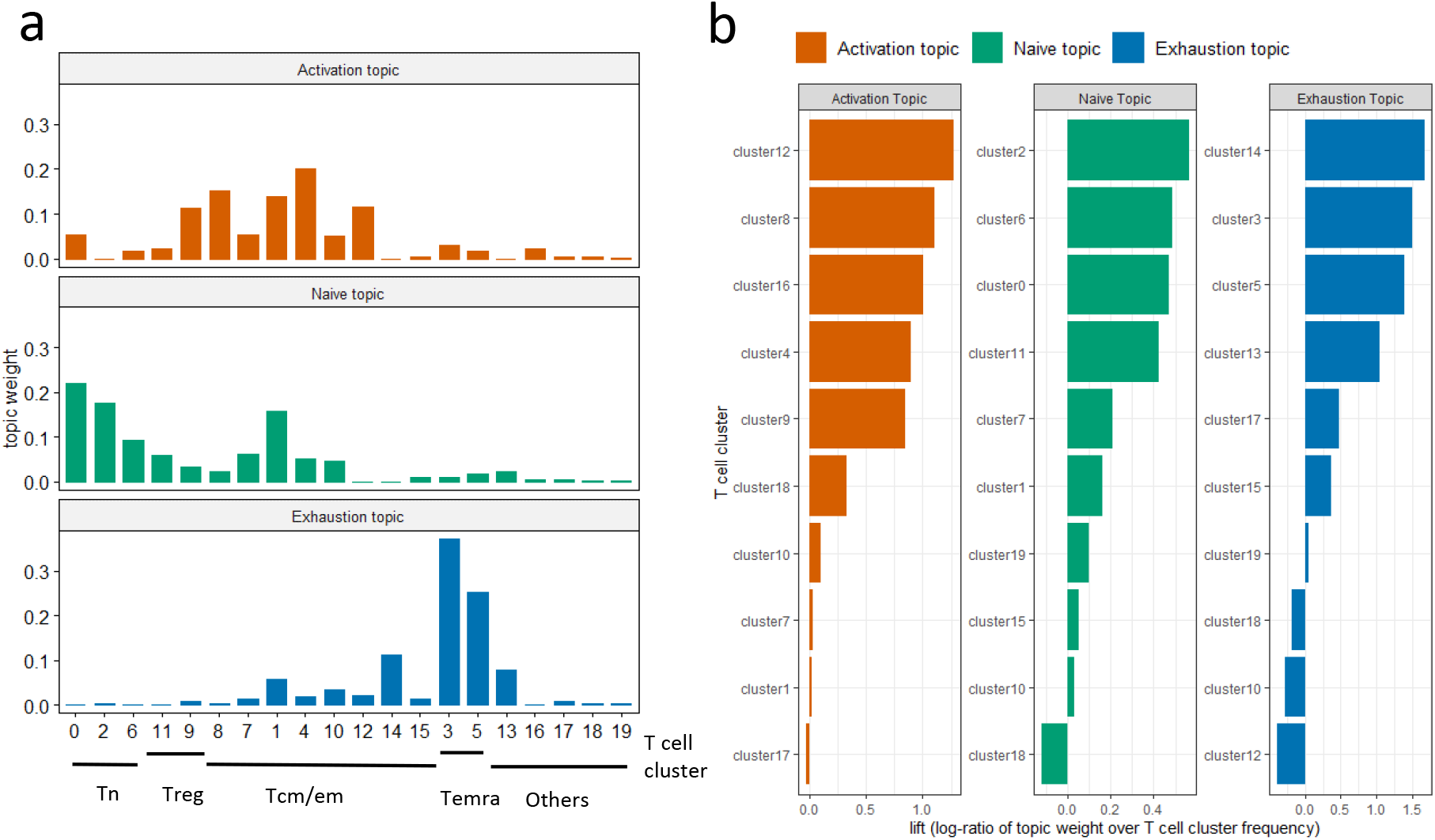
LDA identifies three topics in flow cytometry data. a. Estimated weights (compositions) of clusters *β_k_* in single topics. b. Clusters with the top 10 highest lift for each topic. Clusters with top lift are identified as representative clusters for each topic.

The lift^19^ metric (Fig. 3b, Supplementary Fig. S1), the log ratio of the estimated weight of a T cell cluster v in topic k 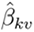 over its empirical frequency, was used to formally rank the importance of individual T cell clusters that characterize each single topic. The biological significance of each topic will be interpreted in the next section, based on their representative clusters with top lift. Each sample can be represented as a mixture of the three topics. The topic-by-sample matrix Θ provides the estimated topic proportions within each sample. Fig. 4a shows the topic fraction across patients and over time. As described earlier, the activation topic mainly captures the expansion of Tcm/em upon treatment. For most patients, the proportion of the activation topic is near zero (dark blue) in pre-treatment samples (week 0). This topic emerges on-treatment as seen by the increase of topic proportions in weeks 3 and 6 samples. At baseline (week 0), most of the patient samples are characterized by a strong presence of the naïve topic. The naïve topic proportion subsequently decreases after ICI treatment as cells transition into more “activated” states. In contrast, a small subgroup of patient samples has a low proportion of the naïve topic, but a high fraction of the exhaustion topic presented at week 0. There is no visible reduction in the exhausted T cell population after ICI treatment.

**Fig. 4:**
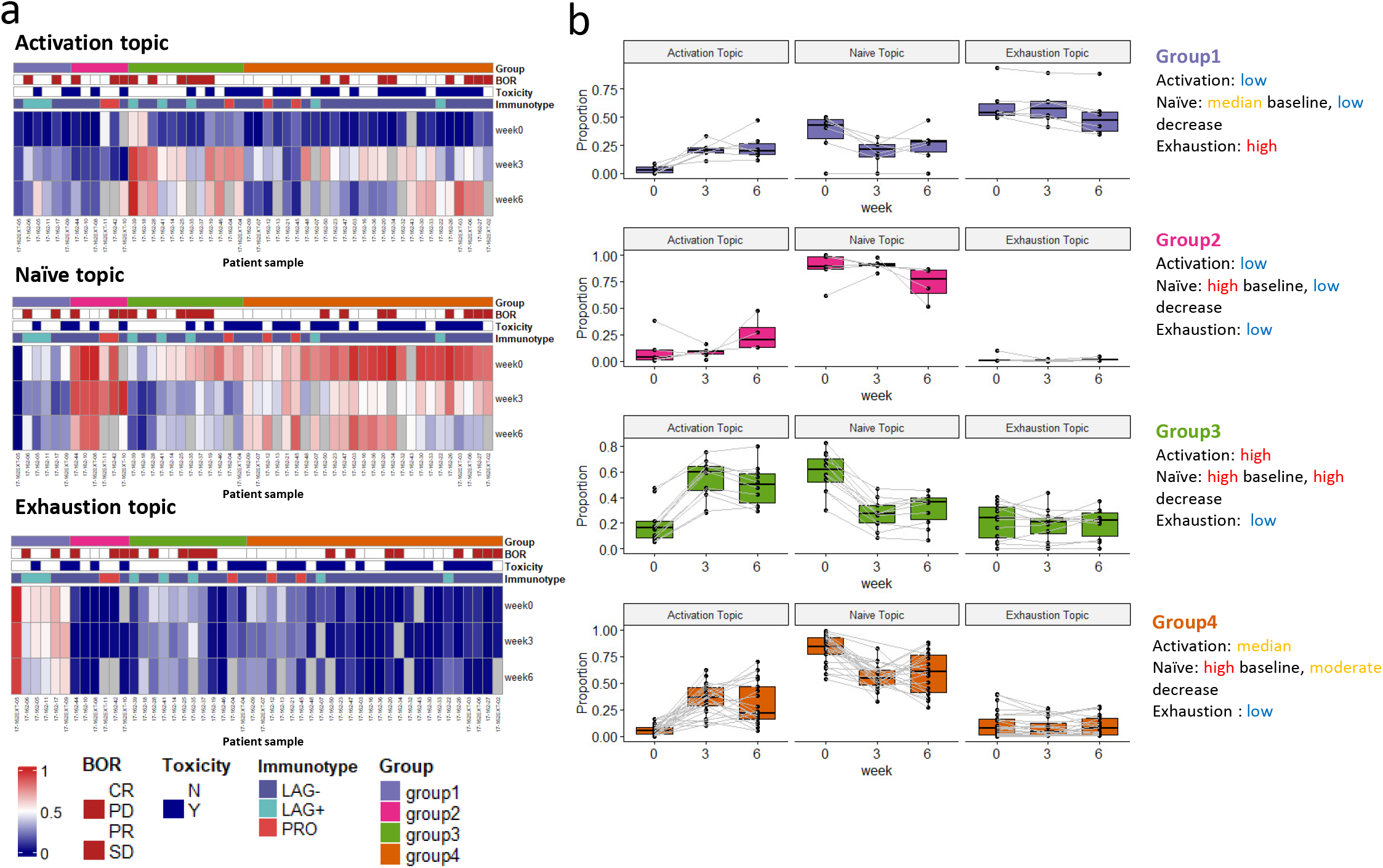
LDA reveals patient subgroups with distinct pharmacodynamics. a. Heatmap showing the sample proportions (*θ_dk_*) for each single topic (patients, n = 50). Patient 17-162-08 has only one sample at week 0, thus it is not included. Missing samples were colored gray in the heatmap. b. Dynamics of sample proportions of the three topics in the four patient subgroups across time.

We identified four patient subgroups by hierarchical clustering on patient topic proportions, while each subgroup exhibits distinct dynamic patterns within the three interpretable topics (Fig. 4b). Patients in groups 1 and 2 both have inferior increases in activation topic. Group 1 has the highest proportion of the exhaustion topic and group 2 has the highest naïve topic across time. Patients in group 3 have the highest increase in the activation topic compared to other groups and are accompanied by the highest decrease in the naïve topic fraction. Group 4 has a high proportion of the naïve topic at week 0 and a moderate increase in the activation topic. Patients in group 4 are more likely to experience severe ICI-related toxicity compared to other groups: 73.1% (19/26) vs 37.5% (9/24) (P = 0.025, Chi-squared test). There is a trend that patients in group 4 have higher response rates: 69.2% (18/26) vs 54.2% (13/24) and better survival outcomes (Supplementary Fig. S2), although not reaching statistical significance.

### Activation topic reveals T cell expansion after ICI treatment

The **activation** topic captures the pattern of T cell expansion in peripheral blood after ICI treatment, as seen by the increase of cells in the representative clusters highlighted in Fig. 5a. The five representative clusters we identified include two CD4 T cell clusters (clusters 8 and 4), one CD8 T cell cluster (cluster 12), one Treg cluster (cluster 9), and one CD4-CD8-T cell cluster (cluster 16) (Fig. 5b). Upon treatment at week 3, the five representative clusters dramatically increased for the entire patient cohort (Fig. 5c), which was captured by the increase in topic proportions (P = 1.3e-33) (Fig. 5d). It might be of clinical interest that most immunological change happens just after the first dose (from baseline to week 3). The comprehensive pharmacodynamics of all 20 clusters are provided in Supplementary Fig. S3-5.

**Fig. 5:**
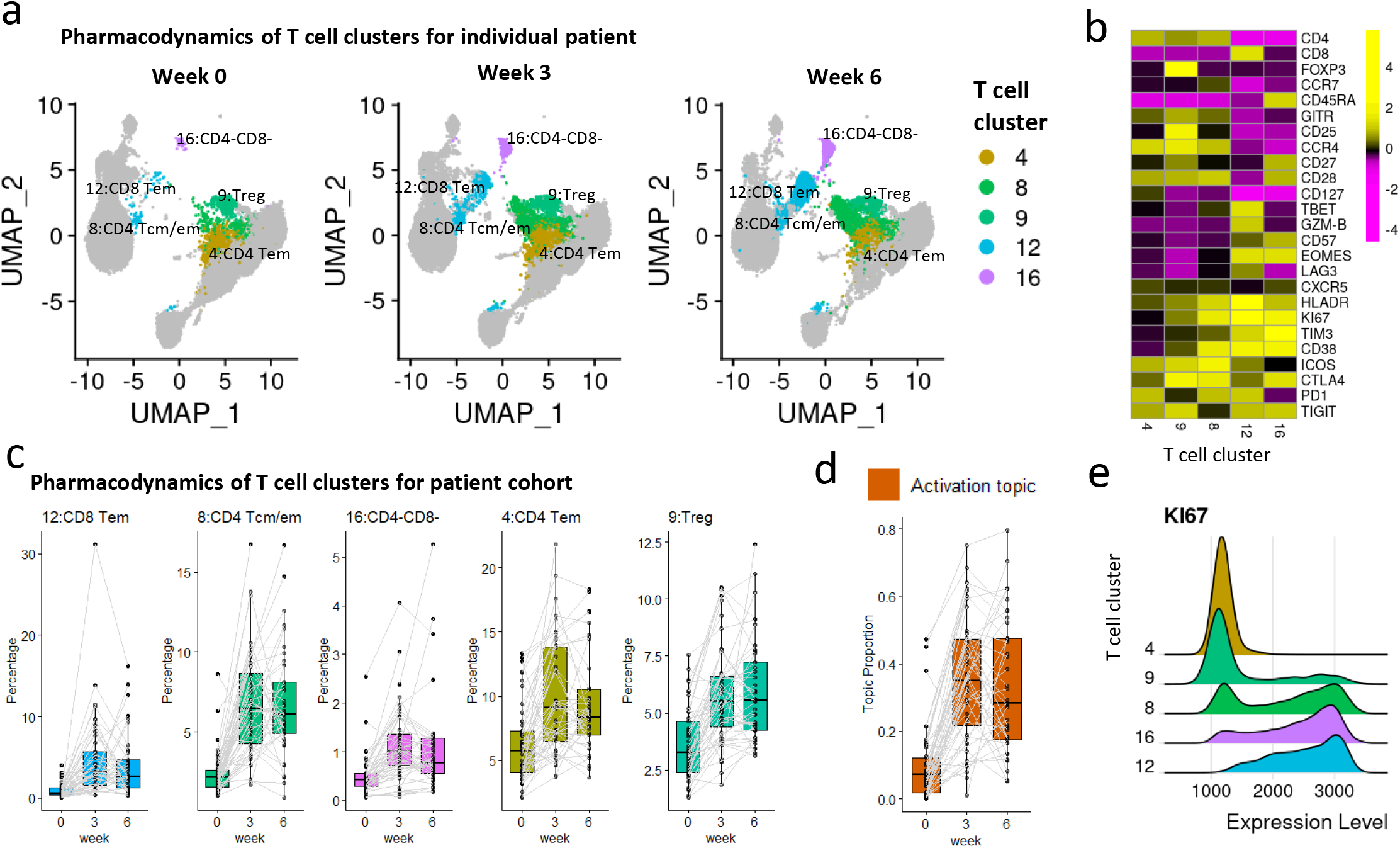
The activation topic. a. UMAP plots of T cells at three time points of patient 17-162-05 (PR, severe irAE), with five representative clusters of the activation topic highlighted. Each UMAP plot contains 20k random-sampled cells from each sample. b. Heatmap showing average marker expression (scaled) of the five representative clusters. c. Relative abundances (percentages of cells in each cluster out of total T cells) of the five representative clusters of the activation topic change over time. The clusters are ordered by lift. d. Activation topic proportions of each individual patient, paired with gray lines. e. Ridge plots of KI67 marker expression over the five representative clusters.

The KI67+ CD8 T cell subset has been established as a T cell reinvigoration biomarker for cancer immunotherapy^9,20,21^. Such a KI67+ CD8 population was independently identified as cluster 12 in our analysis. In addition, cluster 12 also shows high expression of PD1, TIM3, and LAG3 (Fig. 5b), consistent with previous findings that the increase in KI67 expression was most prominent in the PD1+CD8 T cells^9^. In addition to cluster 12, there are two other clusters in our cohort with high KI67 expression: cluster 8 (CD4) and cluster 16 (CD4-CD8-), but distinct in other marker expression profiles (Fig. 5b and 5e). Moreover, we detected an increase in Treg (cluster 9), as observed in another study^9^, with a small fraction of cells expressing KI67 (Fig. 5e). The activation topic presents a novel combination of all these T cell subsets, which can be used as a complex pharmacodynamic index to monitor patients’ immune responses during treatment.

### Naïve topic is associated with ICI-related toxicity

The second topic is a **naïve** topic, with all naïve T cell clusters serving as representative clusters highlighted in Fig. 6a. The four representative clusters we identified include two naïve CD4 clusters (clusters 0 and 2), one naïve CD8 cluster (cluster 6), and one native Treg cluster (cluster 11) (Fig. 6b). The abundances of the four representative clusters, as well as the proportions of the naïve topic, decrease slightly after treatment (P = 5.1e-17 for the difference in proportions across time) (Fig. 6c and 6d), indicating the differentiation of naïve T cells during the immune response. The four representative clusters shared a high level of marker expression in CCR7, CD45RA, and CD27, which are key markers of naïve T cell lineage (Fig. 6b). Interestingly, individuals that experience severe ICI-related toxicity (grade 3-4) have a higher proportion of the naïve topic at baseline week 0 (P = 0.029) (Fig. 6e), while there is no significant difference in changes over time between patients with/without severe toxicity (P = 0.095 for the interaction effect). In contrast, we failed to identify the association between each individual cluster and toxicity (Supplementary Tab. S1), probably due to lack of power after the multiple test correction.

**Fig. 6:**
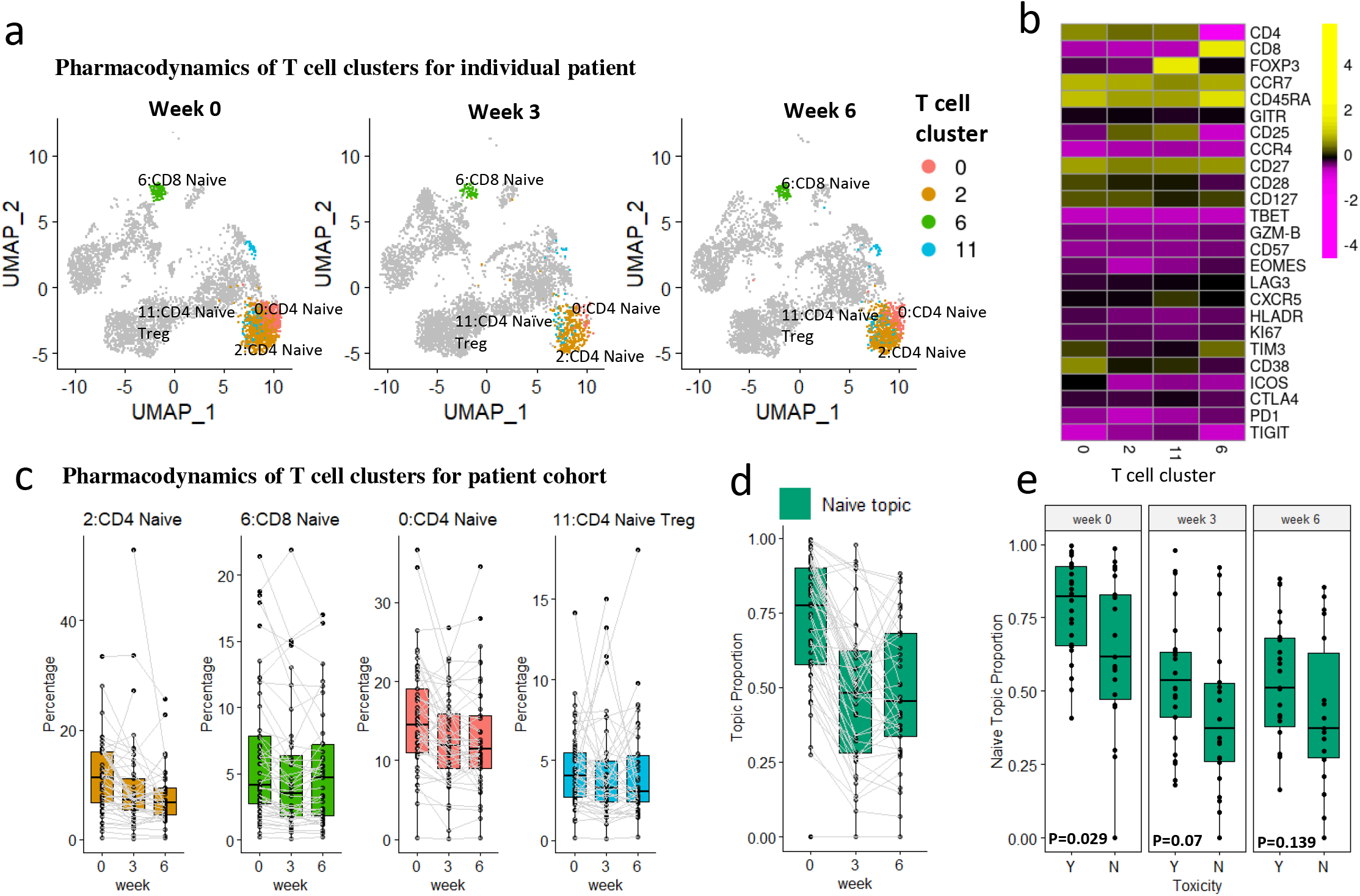
The naïve topic. a. UMAP plots of T cells at three time points of patient 17-162-EXT09 (PR, severe irAE), with four representative clusters of the naïve topic highlighted. Each UMAP plot contains 5k random-sampled cells from each sample. b. The heatmap shows the average marker expression (scaled) of the four representative clusters. c. Relative abundances (percentages of cells in each cluster out of total T cells) of the four representative clusters of the naïve topic change over time. The clusters are ordered by lift. d. Naïve topic proportions of each individual patient, paired with gray lines. e. Sample proportions of the naïve topic between patients experiencing severe/no severe irAE (Y/N). P-values were provided by Wilcoxon rank-sum test for each time point.

### Exhaustion topic is related to LAG+ immunotype

The **exhaustion** topic includes four representative clusters (Fig. 7a): two CD8 Temra clusters (clusters 3 and 5), one CD4 Tem cluster (cluster 14), and one CD4-CD8- cluster (cluster 13). The representative clusters in this topic highly express LAG3, T cell exhaustion marker. Besides LAG3, the four representative clusters also highly express TBET, GZM-B, and EOMES, markers for functional cytotoxic T cells (Fig. 7b). Compared to the other two topics, the topic proportions of the exhaustion topic, as well as the abundances of its four representative clusters, are not significantly changing over time (P = 0.14 for the difference in proportions across time) (Fig. 7c and 7d), but show great heterogeneity in pre-treatment samples (Fig. 4a). For better illustration, we compared pre-treatment samples from two patients (LAG+ vs LAG-immunotype) with four representative clusters highlighted (Fig. 7a). The LAG+ patient sample is dominated by the exhaustion topic (*θ_dk_* = 0.54) while the LAG-patient sample is not (*θ_dk_* = 0.01). We observed substantial differences in abundances of clusters 3, 5, and 14 comparing the two patients.

**Fig. 7:**
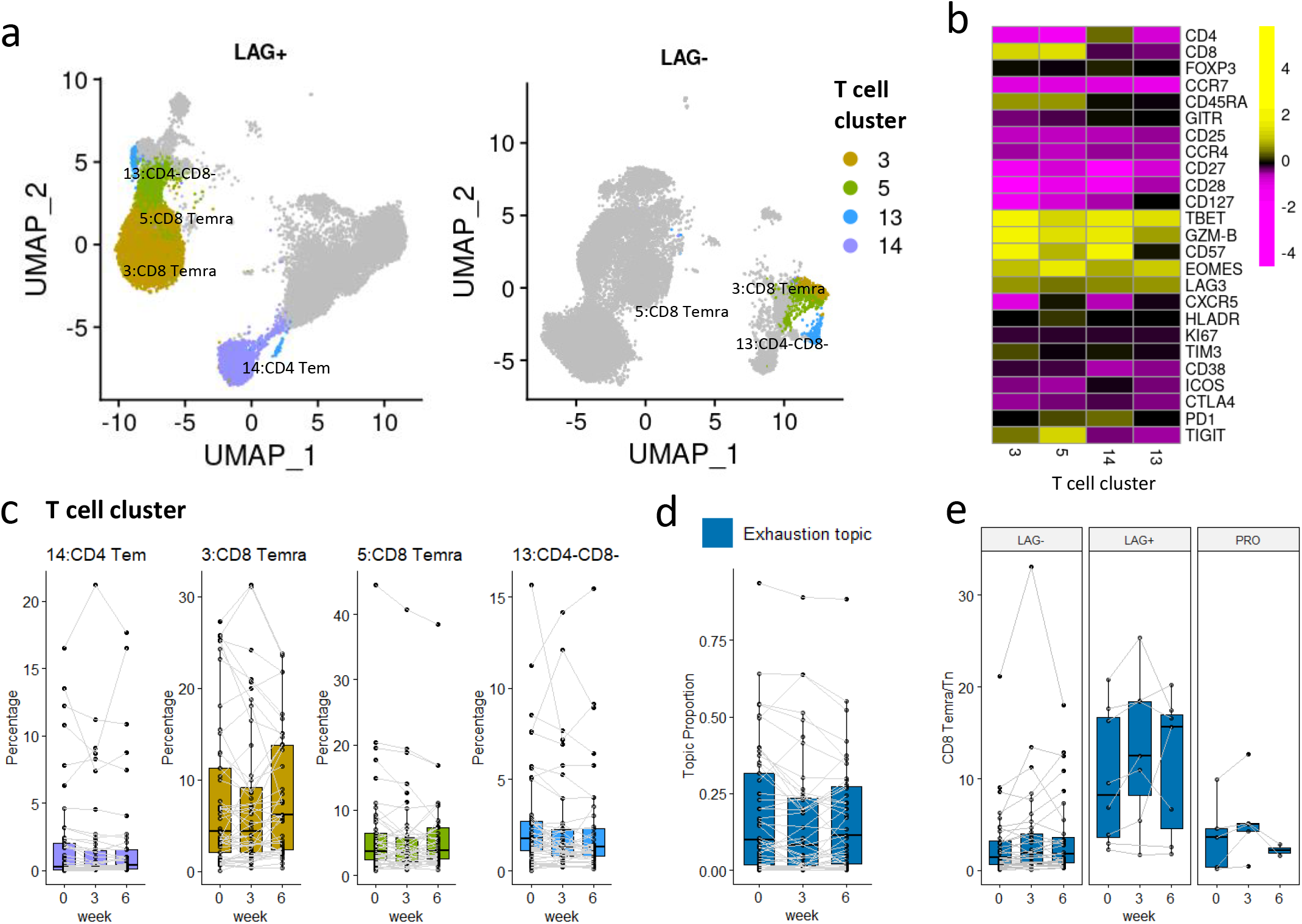
The exhaustion topic. a. UMAP plots of T cells of patients 17-162-05 (PR, severe irAE, LAG+ immunotype) and 17-162-27 (SD, severe irAE, LAG-immunotype) at time point A, each with 20k random-sampled cells. The four representative clusters are highlighted. b. Heatmap of the average marker expression (scaled) of the four representative clusters of the exhaustion topic. c: Relative abundances (percentages of cells in each cluster out of total T cells) of the four representative clusters of the naïve topic change over time. The clusters are ordered by lift. d. Exhaustion topic proportions of each individual patient, paired with gray lines. e. The abundance ratio of CD8 Temra (cluster 3 and 5) to CD8 Tn (cluster 6) across different immunotypes (P = 0.006 for immunotype main effect and P < 0.001 for the interaction effect between time and immunotype). The sample ratios of patient 17-162-EXT05 are extremely high (around ten times the second-highest), and thus are not shown in the boxplot.

The exhaustion topic is highly related to the LAG+ immunotype, which has been linked to poorer clinical outcomes in the earlier study^16^. The previous study classified three immunotypes (LAG-, LAG+, and PRO) on peripheral blood samples using a four-marker classifier (%LAG3+CD8+, %KI67+CD8+, %TIM3+CD8+, %ICOS+CD8+). According to Shen *et al.*, LAG+ patients with high levels of LAG3+CD8+ cells prior to treatment are more likely to have a poor response, particularly with anti-PD-1 regiments^16^. The exhaustion topic provides novel insights into the underlying T cell composition of LAG+/LAG-immunotype. Moreover, we show the ratio of CD8 Temra/Tn (abundances of cluster 3 and cluster 5/ abundance of cluster 6) might be a better biomarker (stable across time and not limited to pre-treatment samples) for distinguishing between LAG- and LAG+immunotype (Fig. 7e), with P = 0.006 for the immunotype main effect and P = 2e-5 for the interaction effect between time and immunotype. This can be explained by the fact that the majority of LAG3+CD8+ cells are Temra cells (in clusters 3 and 5) in pre-treatment samples.

## Discussion

Immune cells are highly heterogeneous, containing a mixture of signals from all unknown ongoing biological processes. Here, we addressed the problem of deciphering hidden structures from longitudinal flow cytometry data in patients treated with ICI. We adopted the LDA model from text analysis and presented a novel computational framework for investigating potentially clinically relevant pharmacodynamical characteristics underlying the data. We demonstrated that LDA is effective in deconvoluting noisy flow cytometry data and can characterize topics that provide novel biological insights. With LDA, T cell subsets can be distilled into topics, which reveal patient subgroups with distinct dynamics.

Our method was inspired by the application of LDA in the longitudinal microbiome analysis^12,13^, where it was able to decipher the temporal changes in microbe composition. Alternative models to monitor dynamics of T cell compositions include the fitness model^22^ from population genetics, and the Lotka-Volterra model (known as the predator-prey model)^23^. However, these models require more time points for model fitting and/or assume no differentiation between cell types. The LDA model on the other hand allows analysis of data from patients with limited time points and was demonstrated to work well on the longitudinal flow cytometry data.

LDA can be further extended and embedded in more complex models for inference. Firstly, incorporating covariates in the topic model could further extend the model application on flow cytometry data, especially under complex experimental design. The Structural Topic Model (STM), for example, allows us to incorporate patient/sample metadata into the model. The metadata can be added as covariates associated with topic prevalence (parameters Θ) or topic content (parameters *B*) with a log link^24^, and a variational Expectation-Maximization algorithm can be implemented for model inference^25^. Secondly, in a setting where long-term monitoring of treatment effects is of interest with a large number of samples collected over time, a dynamic topic model^26^ can be more powerful with a more complex modeling of the temporal relationship across samples. Finally, incorporating additional constraints, e.g. sparsity constraint on cell-type-by-topic matrix *B*, may further improve the efficiency of the model^27^.

The application of LDA is not limited to flow cytometry analysis. For future work, we can further extend LDA to explore the tumor microenvironment in multiplexed imaging data^28^. Spatial information can be incorporated into the model to investigate the tumor and immune cell interactions. Moreover, LDA can also be applied for multi-omics data analysis^29,30^, integrating data from multiple assays to better understand the cancer heterogeneity and predict patient clinical outcomes.

## Methods

### Flow cytometry data

The study includes melanoma patients (n = 51) in a cohort receiving combined immune checkpoint blockade (Anti PD1/CTLA4) therapy from 2017 to 2019 at the Memorial Sloan Kettering Cancer Center in a phase II clinical trial study (NCT03122522)^15^. For each patient, blood samples were collected at three different time points at week 0 (pre-treatment), and at weeks 3 and 6 (post-treatment) after the first dose. Best Overall Response (BOR) [partial response (PR), complete response (CR), stable disease (SD), and progression of disease (PD)], survival, PFS, and toxicity grade [grade 1-2 (N), grade 3-4 (Y)] were determined and reported for each patient. The clinical data for this cohort has been previously described^15^. We also included patient immunotype defined based on the 11-color panel flow cytometry data of pre-treatment samples in our previous study^16^.

The goal of the study is to identify the characteristics of peripheral blood T cells that are related to clinical outcomes (response, toxicity). Flow cytometry with an X50 panel was performed on the collected peripheral blood mononuclear cells (PBMCs) as previously described^31,32^. Our own X50 panel uses a cocktail of antibodies for the following markers: CD45RA-BUV395, CD4-BUV496, ICOS-BUV563, CD25-BUV615, TIM3-BUV661, CD27-BUV737, CD8-BUV805, CD57-BV421, CXCR5-BV480, Live/Dead-FVS510, CD14-BV570, CD19-BV570, CCR4-BV605, CCR7-BV650, HLA-DR-BV711, CD3-BV750, CD28-BV786, PD1-BB515, LAG3-BB660, CD127-BB700, CD38-BB790, TIGIT-PE, EOMES-PE-CF594, CTLA4-PE-Cy5, FOXP3-PE-Cy5.5, GITR-PE-Cy7, TBET-APC, KI67-AF700, GZMB-APC-Fire750. Samples with very poor quality were pre-identified by the flow specialist (M.A.) and were not included in the analysis.

### Pre-gating analysis and quality control

Each Flow Cytometry Standard (FCS) file acquired from the flow cytometry experiments was independently preprocessed using our in-house automated gating pipeline (built with R 4.1.3). The main preprocessing steps include (Supplementary Fig. S6): (1) compensation with matrices exported from FlowJo v10.8.0 software (BD Life Sciences), (2) biexponential transformation on all marker channels with parameters extra negative decades = 0.5, width basis = -30, positive decades = 4.5, (3) quality control via the R package *flowAI* (v1.22.0)^33^, and (4) pre-gating up to CD3+ T cells via the R package *openCyto* (v2.4.0)^34^. The pre-gating strategy is detailed in Supplementary Table S2: a modified version of the T cell gating template originally provided in the *openCyto* R package.

For each marker, we carefully checked the consistency of transformed intensity values across all patient samples, for evaluating the possible batch effects. We downsampled 10k cells from each sample and performed UMAP visualization and clustering analysis on the downsampled data, the same procedure as described in the following clustering analysis section. We visually assessed the UMAP plots and observed no significant batch effect in this cohort. Three samples were excluded in the following analysis due to a lack of cells (<10k cells) for accurate clustering and frequency calculations.

### Clustering analysis

UMAP visualization (min.dist = 0.1) and clustering analysis were performed via *seurat* R package (v4.0)^35^ on pre-gated T cells (CD1419-, CD3+) pooled from all samples. The expression of each marker was scaled to mean 0 and variance 1 before visualization and clustering analysis. Both UMAP and clustering analysis were conducted based on the 26 principal components, using the transformed intensity values of all 27 markers as input. We used the Louvain algorithm, a graph-based clustering method that identifies cell clusters or modules from a Shared-Nearest Neighbor (SNN) graph, a variant of the K-Nearest Neighbor (KNN) graph. We set K = 5 for constructing the SNN graph since it is computationally feasible for over 10 million cells. We ran the clustering algorithms with different resolutions (resolution = 0.5, 0.8, 1.0, 1.2, 1.5, 2, 2.5, 3) and obtained the best clustering result from 10 random starts under each resolution.

We chose the clustering solution under resolution 1.5 based on average Silhouette scores^18^ and manual check. Heatmap was used to show the average (scaled) marker expression of each individual cluster. Clusters of less than 0.1% abundance were not displayed in both UMAP and heatmap to increase the clarity of the figures. We did not include clusters with very low abundance since there is not enough evidence to support that they are real and not generated by technical noises. Moreover, there is no evidence that the low-frequency T cell subpopulations show clinical or biological interests in our analysis. We manually annotated the 20 major T cell clusters (abundance > 0.1%) out of 35 clusters in total. For better visualization, UMAP was rerun for each individual patient with different parameter settings (min.dist = 0.3).

### Latent Dirichlet Allocation

LDA is a generative model that helps to identify hidden structures that explain why some parts of the data are similar. We briefly describe the model and its application to the flow cytometry data below and refer readers to the original paper for more details^14^.

The LDA models the clustered flow cytometry data by considering cells as words, flow samples as documents, and topics as biological profiles or processes. Suppose there are V T cell types (clusters) identified across M samples from S patients. Let *c_dn_* = *ν* for *d* = 1,2, …, *M*, *n* = 1,2,…,*N_d_* represent the nth cell in the *d*th sample classified to the *v*th cell types (clusters). The LDA model assumes each sample has fractional membership across K underlying topics and word *c_dn_* in samples is generated from *z_dn_* th topic, where *z_dn_* ∈ {1,2,…,*K*} are latent variables. In LDA, each sample can be explained by the following generative process (Fig. 1e).

For each sample *d*,

a. Choose sample proportion *θ_d_* ∼ *Dirichlet*(*α*).
b. For each cell *c_dn_* in sample *d*:

1. Choose a topic *z_dn_* ∼ *Multinomial*(*θ_d_*),
2. Choose a cell *c_dn_* conditional on the topic *z_dn_*, *c_dn_*|*z_dn_* ∼ *Multinomial*(*β_zdn_*).

*θ_d_* are mixing proportions of sample *d* over K underlying topics and each topic is characterized as a distribution over V T cell types (clusters), where *β_k_* denote the weights in the *k*th topic over V T cell types (clusters).

In practice, we use the formulation that marginalizes over the *z_dn_*. Setting 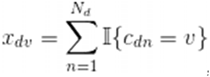, the cell count of the vth cell type in the dth sample, the marginal distribution for each sample d is

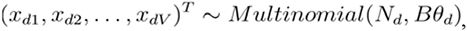

where *B* = (*β_1_*, *β_2_*, …, *β_K_*) denote weights of all topics.

### Model fitting

Gibbs sampling implemented in R package *topicmodels* (v0.2-12)^36^ was used for inferring the two sets of parameters for the LDA model: Θ = (*θ*_1_,*θ*_2,…,_*θ_M_*), a *K*×*M* matrix, and *B* = (*β*_1_,*β*_2,…,_*β_K_*), a *V*×*K* matrix. We used the following setting for Gibbs sampling: iter = 1000, burnin = 1000, thin = 100 (1000 Gibbs sampling draws are made with the first 1000 iterations discarded and then every 100th iteration kept). To evaluate the model reproducibility, we repeated the algorithm ten times and the results of multiple runs are consistent (Supplementary Fig. S7).

The number of topics *K* needs to be selected before running the algorithm and it is a model selection problem. There is no “right” answer to the number of topics that are the most appropriate for data^37^. We failed to select the number of topics with a 10-fold cross-validation, likely a reflection of the size of the dataset (only 138 samples). Thus, we guided the choice of the number of topics based on what is most useful for scientific interpretation. In this study, we set *K* = 3 for the main result in the paper since a larger *K* is less meaningful for only 138 samples.

### Lift statistic

We are interested in representatives, clusters that are primarily associated with a single topic. We use metric *lift*^19^, a popular metric for ranking words within single topics in text analysis, to select representative clusters with the following formula

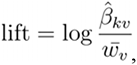

where 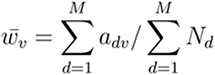 is the empirical frequency of the *v*th T cell type in data, with *a_dv_* being the size of the *v*th T cell type in the *d*th sample. The lift metric gives higher weights to cell types that appear less frequently in other topics.

### Statistical analysis

For each cluster, we also tested its association to clinical outcomes (response, toxicity) and immunotypes via the nonparametric test in *nparLD* R package (v2.1)^38^, which is designed for longitudinal data in factorial experiments. The same method was used to test the association of the ratio (CD8 Temra/Tn), topic proportions to patient clinical outcomes or immunotypes. Only patients with all three time points (n= 37) were included since the package does not support missing data. We included p-values from ANOVA-type tests provided by the *nparLD* R package. For main effects (e.g. immunotypes, response, toxicity) involving only the whole-plot factors, p-values were provided with modified ANOVA-type tests with an adjusted degree of freedom. The Kaplan-Meier method was used for survival estimation and the log-rank test was used for comparisons with the help of survminer R package. Wilcoxon rank-sum test was performed when comparing topic proportions or cluster abundances at each single time point. All p-values from multiple comparisons were adjusted by the Benjamini-Hochberg method with a false discovery rate controlled at 5%.

### Identification of patient subgroups

Patients were grouped by hierarchical clustering (hclust () function in R) on their estimated sample topic proportions Θ. Heatmap was drawn to display the sample topic proportions for each patient, as well as clinical outcomes (response, toxicity) and immunotypes, using the *ComplexHeatmap* R package (v2.10.0)^39^. Boxplot was used to show the dynamics of sample proportions of the three topics within each patient group. One patient (17-162-08) with only one sample at time point A was excluded from the heatmap and the boxplot. Chi-squared tests were performed to test the association between patient subgroups and clinical outcomes (response, toxicity).

## Data Availability

Data file S1 contains all the clinical and correlative data (flow cytometry clusters) analyzed in this manuscript. Additional data for reproducing figures are available in the repository: https://github.com/xiyupeng/topic_modeling.

## Code Availability

Analysis codes to reproduce this work are available in the repository: https://github.com/xiyupeng/topic_modeling.

**The supplementary material pdf includes**

Figs. S1 to S7

Caption for Data File S1

Tables S1 and S2

**Other Supplementary Material for this manuscript includes the following:**

Data File S1

## Acknowledgments

This work is supported in part by MSKCC Society, V foundation, Parker Institute for Cancer Immunotherapy, NIH P30 CA008748, and the MSK-MIND consortium. We thank computational support from MSK-MIND. We thank Jedd. D. Wolchok for help and support on this project. We also thank Nicole Rusk for reviewing and editing the manuscript.

