## Supplementary Material for "Uncovering the hidden structure of dynamic T cell composition in peripheral blood during cancer immunotherapy: a topic modeling approach"

*Corresponding authors

**The PDF file includes:**

Figs. S1 to S7

Tables S1 and S2

Caption for Data File S1

**Other Supplementary Material for this manuscript includes the following:**

Data File S1

**Supplementary Fig. S1: Selection of representative clusters for each topic.**

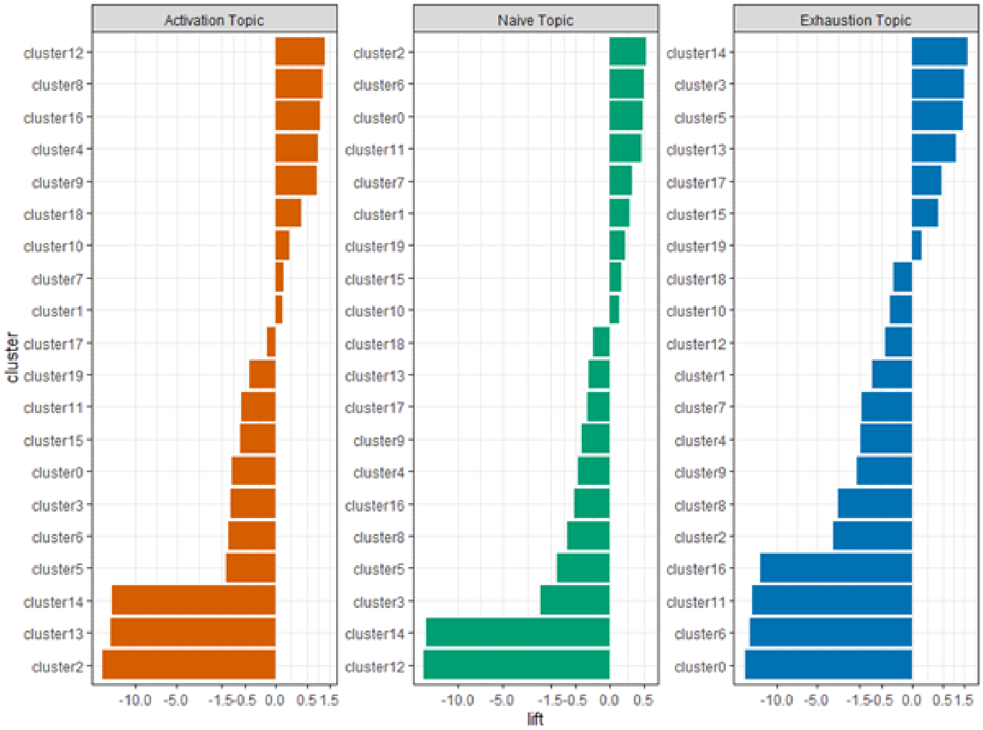


Lift of clusters for each topic, plotted on a signed square root scale. The metric lift gives high weights to clusters that appear less frequently in other topics. Those clusters that have high lift statistics are identified as representatives of single topics.

**Supplementary Fig. S2: Kaplan-Meier analysis of OS and PFS stratified by patient subgroup**

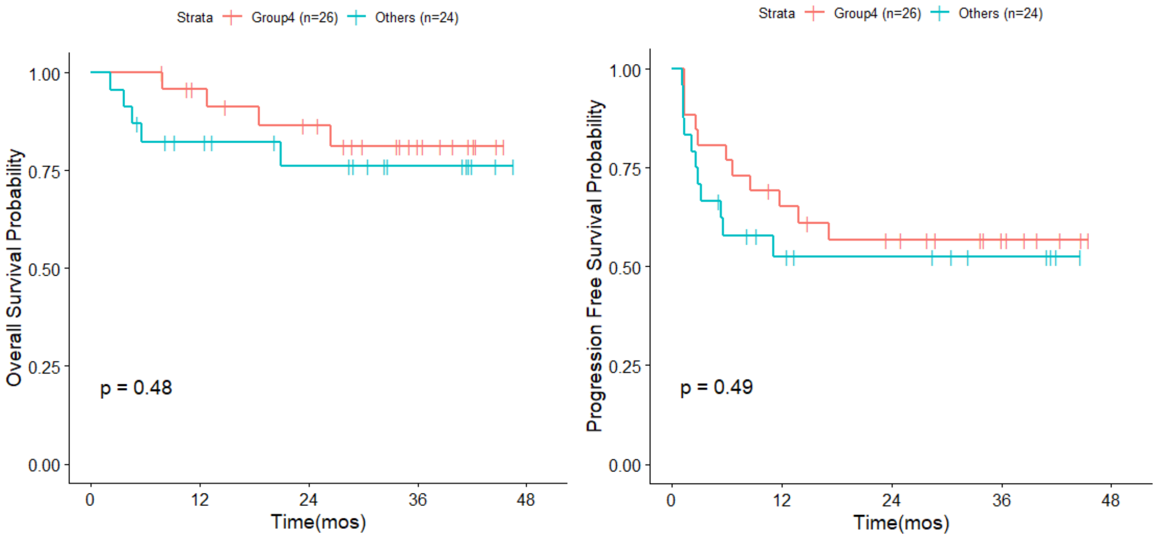

**Supplementary Fig. S3: Pharmacodynamics of single clusters across different immunotypes.**

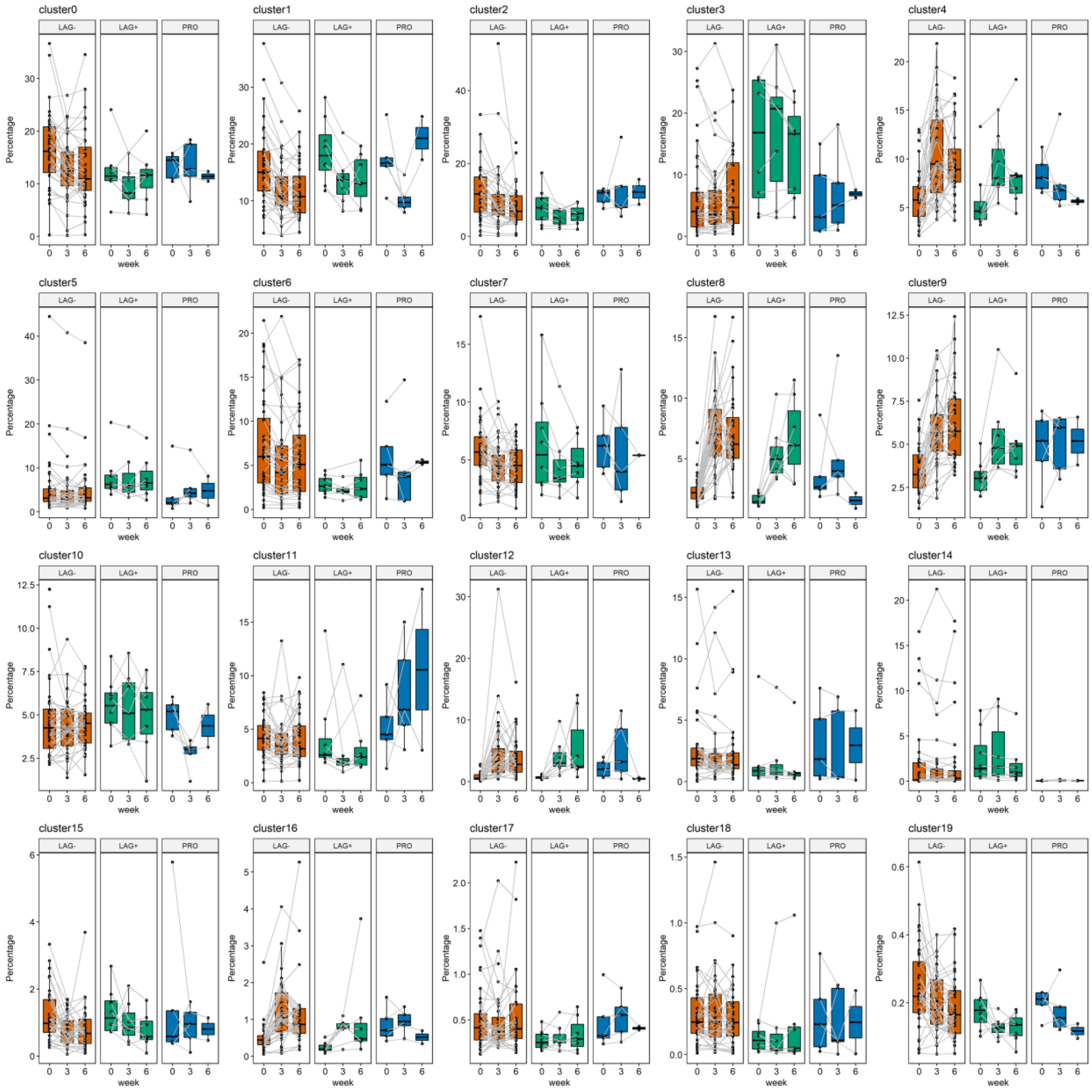

**Supplementary Fig. S4: Pharmacodynamics of single clusters across different responses.**

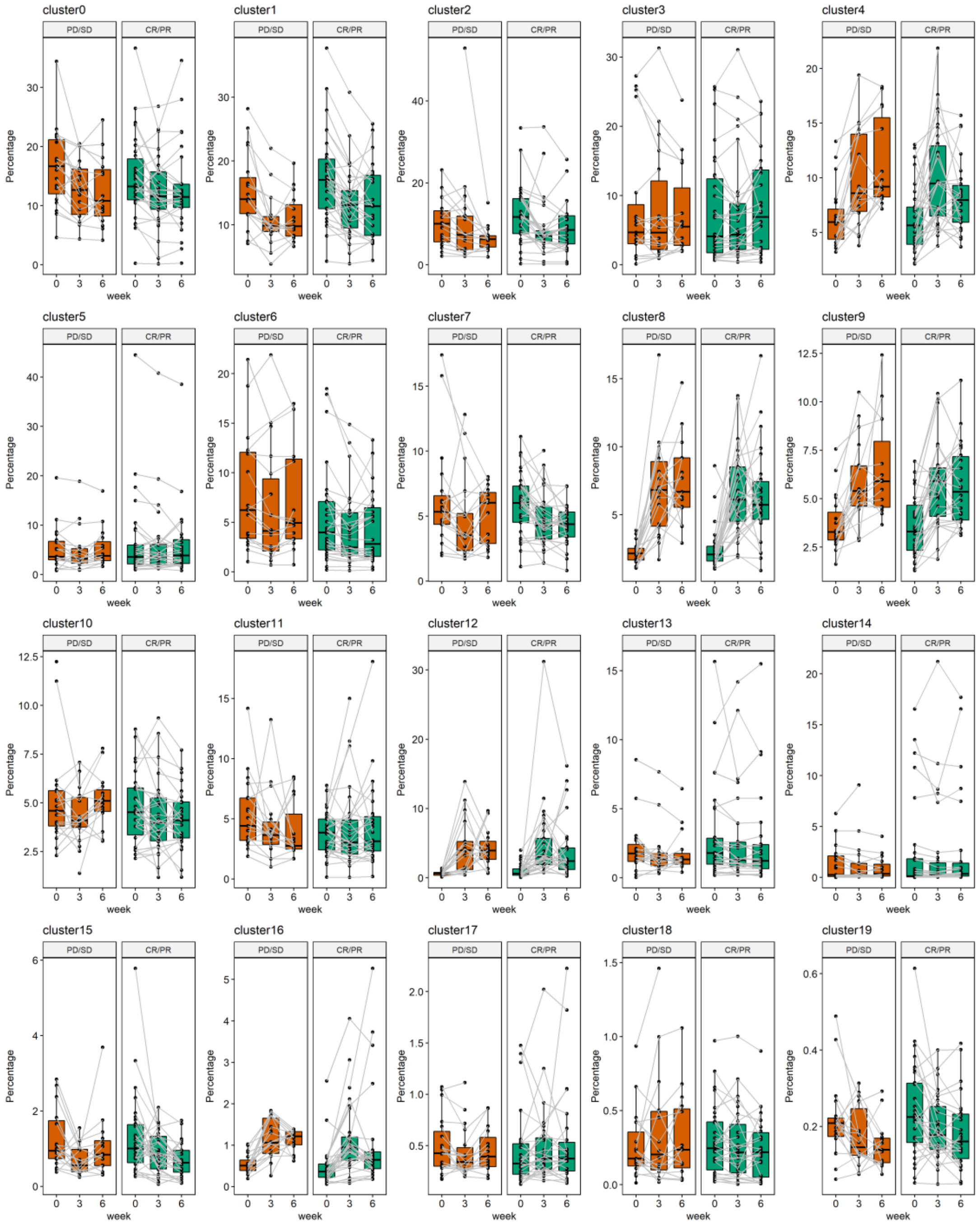

**Supplementary Fig. S5: Pharmacodynamics of single clusters across different levels of toxicity.**

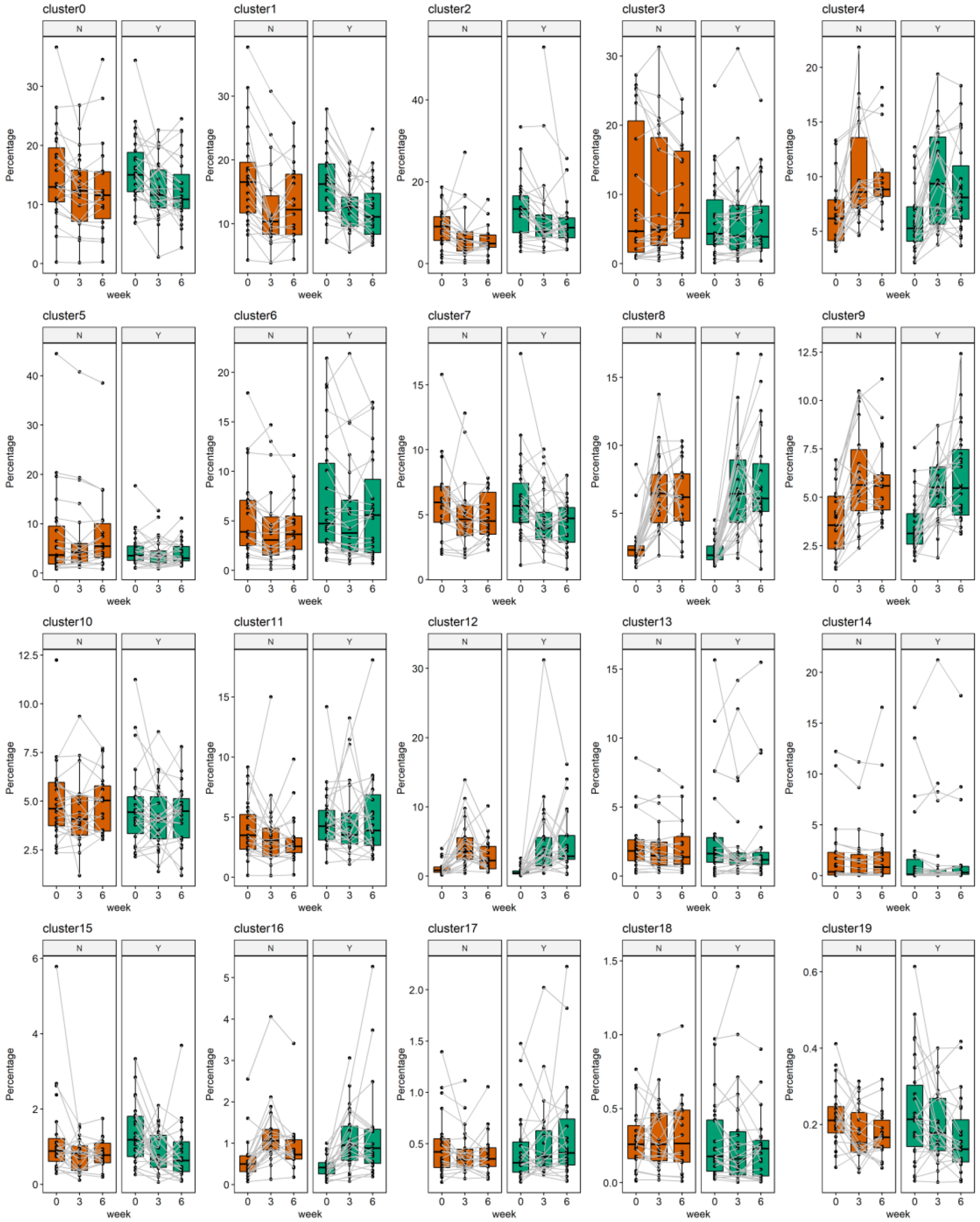

**Supplementary Fig. S6: Pre-gating analysis on flow cytometry data.**

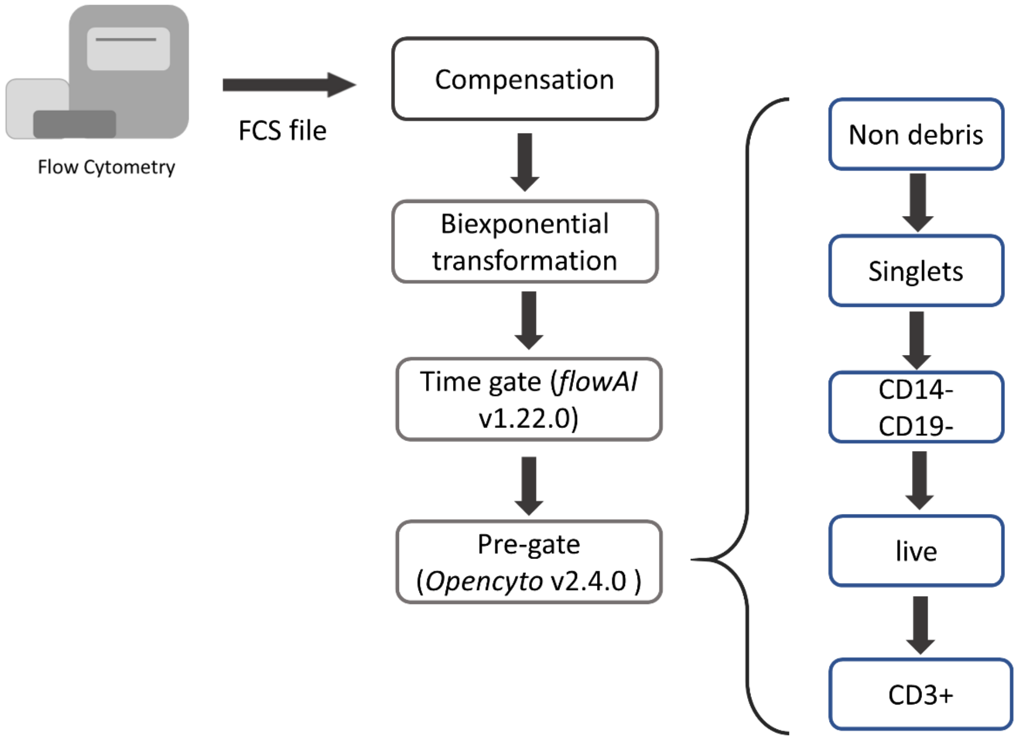

**Supplementary Fig. S7: Estimation of the cell-type-by-topic matrix B by Gibbs Sampling under ten random starts.**

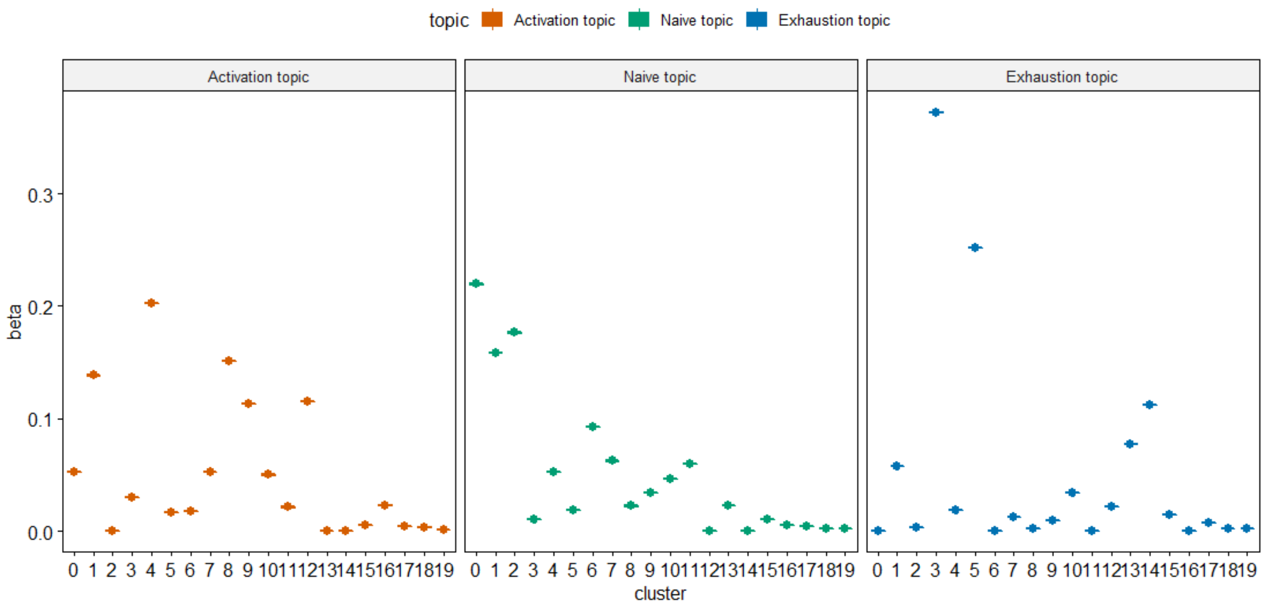

**Supplementary Table. S1. Statistical analysis of single clusters associated with patient clinical outcomes and immunotypes.**

| Cluster | Immunotype | | Response | | Toxicity | |
|---|---|---|---|---|---|---|
| | Immunotype | Interaction with time | Response | Interaction with time | Toxicity | Interaction with time |
| 0 | ns | ns | ns | ns | ns | ns |
| 1 | ns | * | ns | ns | ns | ns |
| 2 | ns | ns | ns | ns | ns | ns |
| 3 | ns | ns | ns | ns | ns | ns |
| 4 | ns | *** | ns | ns | ns | ns |
| 5 | ns | ns | ns | ns | ns | ns |
| 6 | * | * | ns | ns | ns | ns |
| 7 | ns | ns | ns | ns | ns | ns |
| 8 | ns | *** | ns | ns | ns | ns |
| 9 | ns | ns | ns | ns | ns | ns |
| 10 | ns | ns | ns | ns | ns | ns |
| 11 | ns | * | ns | ns | ns | ns |
| 12 | ns | *** | ns | ns | ns | ns |
| 13 | ns | ns | ns | ns | ns | ns |
| 14 | * | ns | ns | ns | ns | ns |
| 15 | ns | ns | ns | ns | ns | ns |
| 16 | ns | * | * | ns | ns | ns |
| 17 | ns | ns | ns | ns | ns | ns |
| 18 | ns | ns | ns | ns | ns | ns |
| 19 | ns | ns | ns | ns | ns | ns |

*** $P < 0.001$; ** $P < 0.01$; * $P < 0.05$; ns, not significant. P-values for the main effect and the interaction effect with time were given by *nparLD* R package, based on patients with all three timepoints (n=37). P-values was adjusted by Benjamini-Hochberg method with a false discovery rate controlled at 5%.

**Supplementary Table. S2. T cell Gating template used in openCyto R package.**

| alias | pop | parent | dims | gating_method | gating_args | collapseDataForGating | groupBy | preprocessing_method | preprocessing_args |
|---|---|---|---|---|---|---|---|---|---|
| nonDebris | + | root | FSC-A | gate_mindensity | | | | | |
| singlets | + | nonDebris | FSC-A,FSC-H | singletGate | | | | | |
| cd14-cd19- | - | singlets | CD14 19 | gate_mindensity | | | | | |
| live | - | cd14-cd19- | L_D | gate_mindensity | | | | | |
| cd3 | + | live | CD3 | gate_mindensity | | TRUE | 4 | | |

In the pre-gating procedure, nonDebris, singlets, CD1419-, live, CD3+ cells were gated in the order described in the gating template, which was used as the input of *openCyto* R package (blank cells are the default to be used as the input). See details in the documentation of *openCyto* R package.

**Supplementary Data File. S1. Sample summary of 51 melanoma patients (xls file attached).**
